# High resolution copy number inference in cancer using short-molecule nanopore sequencing

**DOI:** 10.1101/2020.12.28.424602

**Authors:** Timour Baslan, Sam Kovaka, Fritz J. Sedlazeck, Yanming Zhang, Robert Wappel, Scott W. Lowe, Sara Goodwin, Michael C. Schatz

## Abstract

Genome copy number is an important source of genetic variation in health and disease. In cancer, clinically actionable Copy Number Alterations (CNAs) can be inferred from short-read sequencing data, enabling genomics-based precision oncology. Emerging Nanopore sequencing technologies offer the potential for broader clinical utility, for example in smaller hospitals, due to lower instrument cost, higher portability, and ease of use. Nonetheless, Nanopore sequencing devices are limited in terms of the number of retrievable sequencing reads/molecules compared to short-read sequencing platforms. This represents a challenge for applications that require high read counts such as CNA inference. To address this limitation, we targeted the sequencing of short-length DNA molecules loaded at optimized concentration in an effort to increase sequence read/molecule yield from a single nanopore run. We show that sequencing short DNA molecules reproducibly returns high read counts and allows high quality CNA inference. We demonstrate the clinical relevance of this approach by accurately inferring CNAs in acute myeloid leukemia samples. The data shows that, compared to traditional approaches such as chromosome analysis/cytogenetics, short molecule nanopore sequencing returns more sensitive, accurate copy number information in a cost effective and expeditious manner, including for multiplex samples. Our results provide a framework for the sequencing of relatively short DNA molecules on nanopore devices with applications in research and medicine, that include but are not limited to, CNAs.

## INTRODUCTION

Copy number variation is a common form of genomic variation in humans and has been found to be correlated with a number of pathologies including rare genomic disorders (Vogels and Fryns 2002), neurological diseases (Stefansson et al. 2008; Sebat et al. 2007) and cancer (Shlien and Malkin 2009). In cancer, somatically acquired Copy Number Alterations (hereafter reference to as CNAs) contribute substantially to the remodeling of cancer genomes with diagnostic, prognostic, and therapeutic implications (Döhner et al. 2017; Curtis et al. 2012). Traditionally, CNA information has been retrieved via methods such as chromosome analysis/karyotyping and DNA-FISH (McGowan-Jordan, Simons, and Schmid 2016), which have limitations in sensitivity and/or resolution of analysis. Next generation sequencing addresses these limitations by showing that high resolution CNA information is retrievable from short-read sequencing data using a variety of experimental and computational approaches (Chiang et al. 2009; Baslan et al. 2015; Ha et al. 2012; Kuilman et al. 2015; Xie and Tammi 2009; Wang et al. 2016). These efforts have culminated in the successful implementation of short-read sequencing in the clinic (Zehir et al. 2017; Frampton et al. 2013). However, such applications are still associated with substantial costs and have been largely confined to large, resource rich, clinical centers. Enabling smaller, community-based centers with the ability to retrieve cancer sequence information, in a cost-effective manner, is likely to enhance the quality of health care overall.

New technological advances have paved the path for a newer form of sequencing termed nanopore sequencing (Jain et al. 2016; Goodwin, McPherson, and McCombie 2016). Nanopore sequencing relies on the detection of ionic current fluctuations as DNA molecules translocate through a protein pore (i.e. nanopore) and the subsequent transformation of these signals into nucleotide information (Branton et al. 2008). Nanopore sequencing offers many advantages compared to traditional short-read sequencing technologies, including single-molecule detection, portability, and low instrument cost as well as the ability to retrieve sequence information from large contiguous fragments of DNA, i.e. long read sequencing (Reuter, Spacek, and Snyder 2015). Of these advantages, the ability to sequence long DNA molecules has received the most attention in biomedical research because of its potential to shed light on complex areas of the genome, such as repetitive elements and segmental duplications, areas that remain largely unexplored (Sedlazeck et al. 2018). Further, long read nanopore sequencing has been shown to reveal cryptic genetic variation that is missed by traditional short-read sequencing approaches, accentuating the importance of this emerging sequencing technology in biomedical research and, potentially, clinical practice (Chaisson et al. 2015; Nattestad et al. 2018; Aganezov et al. 2020).

One major shortcoming of nanopore sequencing has been the relatively low yield in terms of the number of distinct sequenced DNA molecules. For example, compared to an Illumina NextSeq machine run, which typically returns hundreds of millions of short-reads, a standard long DNA molecule nanopore run on a MinION device yields an average of approximately one million reads. This limitation has hindered wide-spread utility of nanopore sequencing in research and clinical sequencing applications that require high read counts, especially CNA inference. Here, we apply a simple yet effective experimental approach to address this limitation by loading and sequencing relatively short molecules of DNA and show that it reproducibly returns high read/molecule counts, sufficient for high resolution CNA analysis using the widely studied SK-BR-3 breast cancer cell line as well as five acute myeloid leukemia (AML) samples. Our approach provides a foundation for future studies where real-time, point of care sequencing applications in cancer, and other diseases, can be done with minimal infrastructure in clinical centers, large and small alike.

## MATERIALS AND METHODS

### Samples included in study, DNA extraction, and sequencing library preparation

To develop the short molecule sequencing approach, we first experimentally tested DNA library preparation conditions and sequencing on DNA retrieved from a widely studied breast cancer cell line, SK-BR-3, which we and others have used as a reference for CNA method development (Baslan et al. 2015; Garvin et al. 2015; Navin et al. 2011). For nanopore sequencing library preparation, SK-BR-3 cell line DNA was fragmented to approximately 500bp using the Covaris Instrument (Woburn, MA). 300ng of sonicated material was A-tailed and end repaired using the NEB A-tailing end repair module. Adaptor ligation for nanopore sequencing was performed using the SQK-LSK-109 kit from Oxford Nanopore. Three different DNA library loading masses were tested to determine optimal loading condition; 15ng, 45ng and 90ng. 15ng is equivalent to 50 fmol of dsDNA; the maximum, manufacturer suggested mass gDNA for loading. We hypothesized that a larger mass of DNA could improve throughput for smaller fragments, thus 45ng (3X the suggested molar mass) and 90ng (6x the molar mass) were also tested. Each aliquot of library was loaded onto a R9.4.1 MinION flow cell and allowed to run for 48 hours on a GridION. For clinical implementation feasibility studies, we relied on sequencing DNA retrieved from acute myeloid leukemia (AML) samples. Five samples were considered for MinION sequencing, two that were cytogenetically classified as normal karyotype and three classified as complex karyotype (CK-AML), with the latter known to confer a dismal prognosis in AML disease. We also sequenced two additional AML samples, one complex and one normal, using the low throughput Flongle. DNA was extracted from bone marrow leukemic blasts using a Qiagen AllPrep DNA/RNA Mini Kit following recommended manufacturer’s protocols. Purified DNA was similarly processed for nanopore sequencing as per final experimental conditions established using SK-BR-3 DNA. For multiplex nanopore sequencing, DNA was prepared as described above with the following exception: after the initial A-tailing and end repair step, DNA was ligated to a PCR based barcoding adapter (EXP-PCB096) for unique indexing. Five barcoded samples were then pooled in an equimolar fashion, with a total mass of 300ng between all five samples. The pooled samples were processed with the SQK-LSK-109 kit. Samples were base called with the on-board GPU system with Guppy v3.1. The research involving human subjects, specifically the AML samples, was approved by the authors Institutional Review Board (MSKCC IRB).

For each analyzed sample (SK-BR-3 as well as AML samples), a matching short-read Illumina sequencing library was constructed and sequenced on a HiSeq device using standard Illumina protocols (San Diego, CA).

### Nanopore sequence read base calling, processing, and analysis

For each sequencing run we computed read length distributions, read counts over time, channel lifetimes, molecule residence time (i.e. the time duration for an actively sequenced read), pore vacancy time (i.e. the time between reads in a given channel), and channel activity. All summary statistics provided in **Table 1** based on the “sequencing summary” files output by the Guppy basecaller. These summary files include the time that each read started and finished sequencing, which channel reads came from, read lengths, and quality scores. Read counts were computed using a simple cumulative sum over the course of each run. “Relative Reads per Channel” was computed by dividing the reads per channel by the number of channels on the device (512 for MinION, 126 for Flongle). Channel lifetimes were estimated by the time that each channel finished sequencing its final read. The distribution of vacancy time between reads was computed by subtracting the start and end times of consecutive reads, excluding reads which were not sequenced by the same pore within a channel. The channel activity plots were generated by dividing each channel into five minute windows and coloring a window black if the channel sequenced any length of a read during that time.

**Table 1.** Annotations and sequencing statistics for samples run on a MinION device. AML = Acute Myeloid Leukemia, ME = Molar Equivalent. All AML samples were run at 3 ME.

| Sample/<br>Type | Sample.ID | Read<br>Count | Yield<br>(Mbp) | Passed<br>(%) | Q-Score<br>Mean | Read- Length<br>Mean | Read<br>N50 | Minutes to<br>250k reads |
| --- | --- | --- | --- | --- | --- | --- | --- | --- |
| Cancer cell line<br>SKBR3 | SKBR3 - Long | 899,425 | 8,360.86 | 84.94 | 10.17 | 9,295.79 | 14,301 | 291.93 |
| Cancer cell line<br>SKBR3 | SKBR3 - Short<br>(1 ME) | 6,332,668 | 3,024.80 | 86.42 | 10.54 | 477.65 | 538 | 16.50 |
| Cancer cell line<br>SKBR3 | SKBR3 - Short<br>(3 ME) | 1,465,247 | 789.33 | 89.61 | 10.89 | 538.70 | 637 | 17.07 |
| Cancer cell line<br>SKBR3 | SKBR3 - Short<br>(6 ME) | 699,009 | 416.13 | 88.69 | 10.73 | 595.32 | 732 | 19.35 |
| Leukemia<br>(AML-BM) | AML1 | 3,797,811 | 1,670.22 | 84.97 | 9.09 | 439.79 | 501 | 15.13 |
| Leukemia<br>(AML-BM) | AML2 | 6,263,490 | 2,886.35 | 82.82 | 8.95 | 460.82 | 519 | 15.53 |
| Leukemia<br>(AML-BM) | AML3 | 3,935,726 | 1,753.57 | 86.68 | 9.29 | 445.55 | 488 | 15.58 |
| Leukemia<br>(AML-BM) | AML4 | 3,935,726 | 1,072.69 | 91.29 | 9.72 | 463.96 | 503 | 16.59 |
| Leukemia<br>(AML-BM) | AML5 | 6,065,542 | 2,510.64 | 81.17 | 8.69 | 413.92 | 461 | 16.42 |

### Copy number inference from short read and short molecule sequencing data

Nanopore sequencing data was mapped to human reference genome hg19 using Minimap2 (Li 2018) software with the following parameters: -ax map-ont −t 15. Uniquely mapped reads were subsequently filtered using Samtools (Li et al. 2009) with the following parameters: -Sbu -q 20 -F 0×904. Sorted reads were indexed and counted in genomic bins using the Varbin algorithm at two different resolutions; 5 thousand and 20 thousand, 5k and 20k respectively, derived by dividing the genome into the aforementioned bins respectively (Navin et al. 2011; Baslan et al. 2012). Normalized bin counts were subsequently segmented using Circular Binary Segmentation (Olshen et al. 2004) and transformed to absolute copy number values using a least square fitting algorithm (Baslan et al. 2015; Garvin et al. 2015). Short read Illumina sequencing data was processed as previously described (Baslan et al. 2015).

### AML sample chromosomal analysis

Fresh bone marrow aspirates were used for conventional chromosome analysis following standard protocol, without mitogen stimulation in short term culturing. At least 20 metaphase cells were analyzed, and karyotype was described according to the international system of human chromosome nomenclature (McGowan-Jordan, Simons, and Schmid 2016). Independent numerical and structural chromosomal abnormalities, such as gain, loss, deletion, addition, duplication, balanced or unbalanced translocations, inversion and derivative chromosomes, as well as marker chromosome, ring chromosome were annotated in defining complex karyotypes, i.e., three or more, clonal heterogeneity, and monosomal karyotype

### DNA-FISH validation of identified copy number alterations in AML

FISH analysis was performed on bone marrow or peripheral blood pellets from cytogenetic analysis in selected samples, following standard protocols. Various commercial FISH probes, specific for myeloid neoplasia, such as for deletion or loss of chromosomes 5, 7, 17, gain of chromosome 8, and for MLL/KMT2A (11q23) translocations, EVI1 (3q26.2), and other translocations, were used as appropriate. All probes were purchased from several companies, such as Abbott Molecular (Des Plaines, IL), Metasystems (Newton, MA). At least 200 cells were analyzed, and results were described according to (McGowan-Jordan, Simons, and Schmid 2016). In all samples, correlation of FISH results and chromosome analysis findings were also compared.

## RESULTS

### Sequencing short DNA molecules on a nanopore device yields high read counts and enables accurate copy number profiling

The number of reads/molecules returned in a standard long read MinION run is limited by (1) the design of the nanopore array, especially the overall number of channels and the overall number of active pores; (2) the molecule kinetics of DNA fragments docking to the nanopore, measured by the duration between the end of one read and the beginning of the next within the same pore (the vacancy time); and (3) the residence time of a molecule translocating through a given pore. Consequently, we hypothesized that loading DNA molecules that are shorter in length (e.g. 400bp in median length compared to a standard 10kb) would decrease the residence time of each DNA molecule in a pore and thus facilitate, over a period of time, the translocation of more DNA molecules and the retrieval of higher read/molecule counts.

To experimentally test our hypothesis, we sonicated DNA purified from the SK-BR-3 breast cancer cell line to an average length of 400 bps and prepared a nanopore sequencing library using standard protocols (**Methods**). For short read/molecule sequencing on a R9.4.1 MinION flowcell, we decided to load libraries at concentrations of 1, 3, and 6X the Molar Equivalent (ME) of standard loading conditions (**Methods**), henceforth referred to as Short-1ME, Short-3ME, and Short-6ME respectively. Varying the loading conditions we performed to ascertain the relationship between number of molecules loaded and subsequently sequenced in each experiment. As a control, a long read sequencing library from the same SK-BR-3 DNA generated and sequenced using standard protocols (**Methods**).

Sequencing the long molecule nanopore library resulted in an expected yield of approximately 1 million molecules. By contrast, loading the Short-1ME sequencing library resulted in roughly 6 million sequenced molecules, a six-fold increase (**Fig 1A**). Interestingly, while the number of active sequencing channels decayed with a similar trend for both sequencing conditions (**Fig 1B**), the increased read yield in the Short-1ME library was largely attributed to an accelerated return of sequenced molecules over the first 12 hours (**Fig 1A, S1A Fig)**. We attribute the increase in sequenced molecule return primarily to reduced residency and vacancy time. Residency time is directly proportional to the length of the molecules and the speed of sequencing, for example approximately 1 second for a 400 bp molecule compared to 22 seconds for a 10 kbp molecule when sequencing at 450 bp per second. Further, vacancy time is also shorter when sequencing short molecules, regardless of loading concentration **(Fig. 1C)**. Short-1ME sequencing reads were of similar quality to long reads when comparing sequencing metrics such as Pass Filter (86% Short-1ME vs. 85% Long) and mean Q-Score values (10.54 Short 1ME vs. 10.17 Long) (**Table 1**). The average length of the sequenced DNA molecules was confirmed to be roughly 500bps (**S1B Fig)**.

**Fig1.**
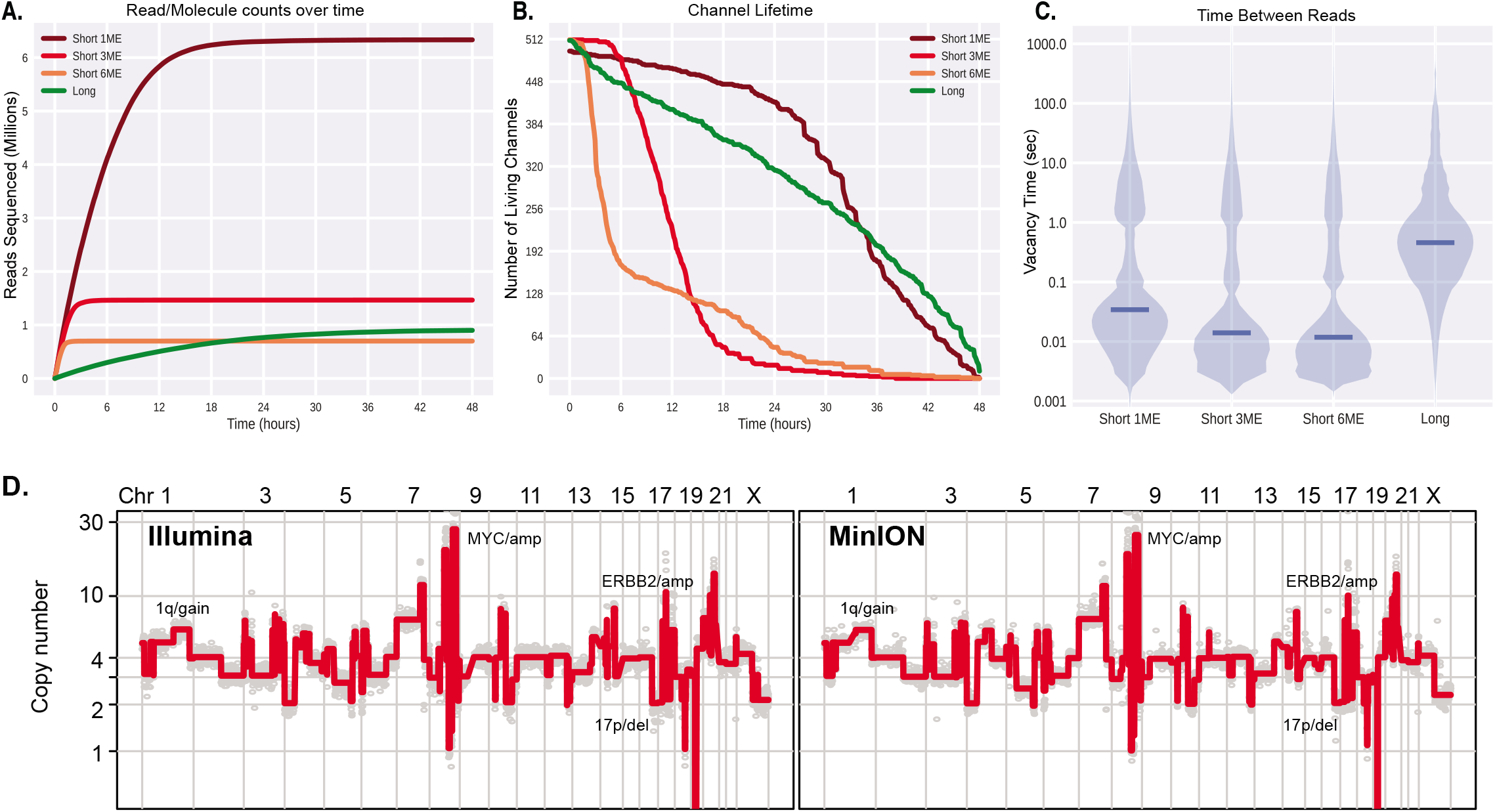
Sequencing short molecules on a nanopore device yields high read counts and enables accurate copy number profiling. (A) Cumulative number of reads/molecules sequenced over time throughout each SK-BR-3 run (ME = Molar Equivalent). (B) Total number of living channels over time throughout each run as measured by the time that the last read was sequenced by each channel. Note that a channel may not produce a read for several hours but still be considered alive by this definition. (C) Distributions of timing between discrete DNA molecules entry/exit from a given channel – i.e. vacancy time. (D) Genome-wide copy number profiles of SK-BR-3 sequenced using short molecule nanopore (right panel) and short-read Illumina (left) data. Profiles at plotted when dividing the genomes in 5 thousand bins (i.e. 5k bins). Examples of detected CNAs are annotated on the profiles.

Importantly, when processed for copy number inference using previously developed algorithms (Baslan et al. 2015; Garvin et al. 2015) that relate read depth (i.e. molecules sequenced at a given genome locus) to genome copy number, we find that the data produces highly accurate copy number information at various resolutions compared to information retrieved from standard Illumina short-read sequencing data (**Methods**) (**Fig 1D and S2A Fig)**. These included: (1) focal amplifications at the *MYC* and *ERBB2* genes, (2) gain on 1q, and (3) deletion on 17p, all CNAs shown to be associated with a poor prognosis across a range of human tumors. For both the Illumina and Short-1ME sequencing data we corrected for GC nucleotide content biases using our standard LOWESS smoothing algorithm (Baslan et al. 2015) (**S2B Fig)**.

Short-3ME and Short-6ME libraries yielded 11 and 6 times more sequenced reads respectively relative to long reads over the first two hours, however the sequencing yield pattern decayed rapidly (**Fig 1A)**. This pattern was associated with a diminishing number of active channels over time as well as vacancy time relative to the Short-1ME sequencing experiment (**Fig 1 A,B, and C)**. As the quality of the data is similar to that from long and Short-1ME data **(Table 1, S1B Fig)**, we hypothesize that this decline may be a consequence of the mechanics of nanopore/electrical circuitry and/or limiting reagent availability to drive the translocation of DNA molecules through the MinION nanopore channels.

Overall, these data show that loading short DNA molecules, at an appropriate concentration, to a MinION nanopore device can yield up to 6 million sequenced molecules in a single run, data which can be used to infer CNAs with high accuracy.

### Short molecule sequencing on a MinION device allows accurate, high resolution copy number inference in a clinically relevant setting

To assess the potential utility of short molecule sequencing on MinION devices in retrieving clinically actionable and relevant information, we turned to the analysis of Acute Myeloid Leukemia (AML) genomes where extensive evidence exists on the prognostic importance of CNAs (i.e. karyotypes/cytogenetics). Five samples were sequenced, two that were cytogenetically classified as normal karyotype and three classified as complex karyotype (CK-AML), with the latter known to confer a dismal prognosis in AML disease. All AML samples were sequenced using Short-1ME loading conditions. In parallel, all samples were sequenced on an Illumina machine for comparative analysis.

All AML samples sequenced using Short-1ME loading conditions yielded expected high read counts (∼4-6 million reads per sample) (**Table 1)**. Further, all samples displayed similar high-quality sequencing statistics, such as Pass Filter and mean Q-Scores, as with the Short1-ME SK-BR-3 data (**Table 1)**. When processed for copy number inference, we find that the nanopore Short-1ME data yields largely similar genome-wide profiles and CNAs with Pearson correlation coefficients above 0.98 across all samples when compared to Illumina data (**Fig 2A, B, C and S3 Fig)**. For example, for sample AML2, nanopore sequence data inferred CNAs that were highly correlated to those inferred from Illumina sequencing data (Pearson correlation values > 0.99) with large deletions on chromosomes 5 and 7 (over 10MBs) as well as complex re-arrangements on chromosome 11 resulting in *MLL1/KMT2A* gene gains identified. Further, identified alterations shared largely identical breakpoint positions in both datasets **(Fig 2B, C, and D)**. Moreover, focal alterations on chromosome arms 12p and 16p were also identified in both nanopore and Illumina sequencing data and corroborated with standard cytogenetic analysis (**Fig 2E and S3 Fig)**.

**Fig2.**
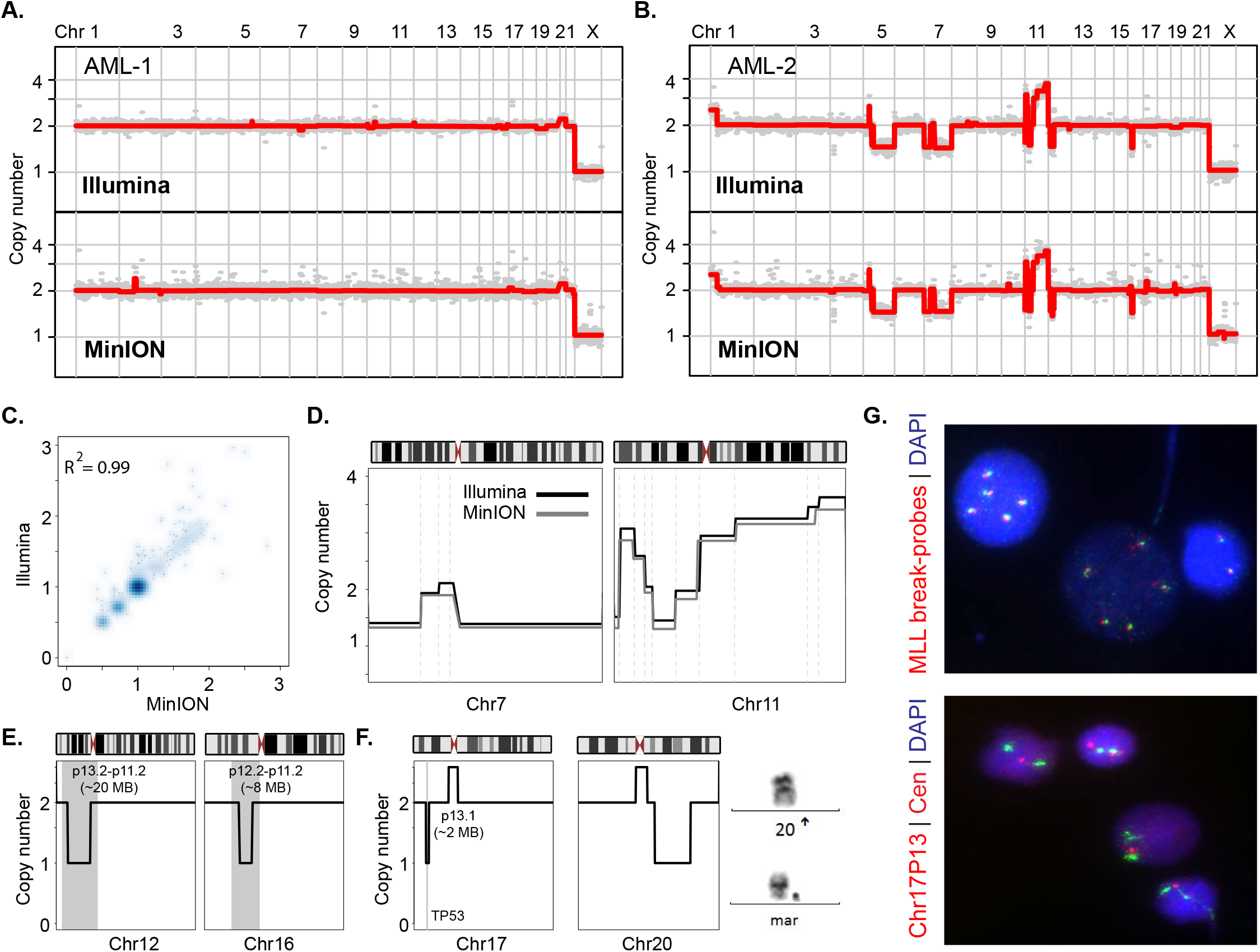
Short molecule sequencing on a MinION nanopore device yields accurate high resolution copy number information in a clinically relevant setting. (A) and (B) Genome-wide copy number profiles of a normal and a complex karyotype sample sequenced using short molecules on a MinION (lower panel) and short-read on an Illumina instrument (upper panel). (C) Density scatter correlation plot of normalized bin read counts (proportional to copy number) from MinION and Illumina sequencing data for leukemic sample AML-2. Pearson correlation value is provided. (D) Zoom in views of chromosome 7 (left panel) with complex rearrangement resulting in loss of both short and long arms and chromosome 11 (right panel) with gains of the long arm at various level in complex karyotype sample AML-2 using Nanopore (gray line) and Illumina (black line) sequencing data. Dashed gray vertical lines illustrate alteration breakpoints. (E and F) Zoom in views of chromosomal deletions on chromosomes 12p, 16p, 17p and 20q. Right panel in (F) illustrates chromosome images derived from cytogenetic analysis showing a normal chromosome 20 (top) and a marker chromosome (bottom) which was redefined as a derivative chromosome 20 with deletion of the long arm based on this study (G) DNA-FISH based validation of selected CNA alterations. Gain of the MLL gene on chromosome cytoband 11q23 (top panel), and the focal 17p loss, encompassing the TP53 gene (bottom panel)

In multiple cases, nanopore sequencing data resulted in more accurate copy number information compared to standard cytogenetic chromosome analysis. For example, for sample AML5, cytogenetically reported monosomy (i.e. loss of an entire chromosome) for chromosome 20 was shown in both Illumina and nanopore sequencing data to be inaccurate, most likely the result of incorporation of chromosome 20 genetic material into marker chromosomes (**Fig 2F)**. Likewise, copy number analysis at higher resolution, afforded as a direct consequence of the increased read counts, identified a focal deletion on the short arm of chromosome 17 (17p) encompassing the *TP53* gene in the sequencing data (**Fig 2F**). Selected alterations (*MLL* and *TP53*) were validated using DNA-FISH analysis in their respective samples (**Fig 2G)**.

Together, these results show that short molecule sequencing on a MinION device reproducibly yields high read/molecule counts that are of high quality. Further, when processed informatically, the data results in highly accurate copy number profiles that are concordant with both matching Illumina data and standard cytogenetic results. The data also show that short molecule nanopore sequencing returns cryptic CNA information that is either missed or mis-classified in standard cytogenetic, chromosome-based analyses.

### High read counts enable multiplex sequencing and copy number inference of AML samples on a MinION device

Sequencing-based copy number inference tools can rely on sparse, low coverage sequencing to relate sequencing depth in genomic intervals to copy number (Baslan et al. 2015; Garvin et al. 2015). Furthermore, sequencing depth can be adjusted to accommodate varying resolution of analysis, typically targeting an average of 30 to 50 reads per genomic bin for a reliable CNA inference (Baslan et al. 2015; Garvin et al. 2015). Given the comparatively high number of sequencing molecules retrieved by sequencing Short-1ME nanopore libraries, multiplexing of different samples on the same MinION run is theoretically possible. Multiplexing would lower the costs associated with retrieving sequencing-based copy number information as well as provide flexibility in terms of sample processing, both issues that are relevant from a clinical implementation perspective. To this end, we generated uniquely indexed short molecule nanopore sequencing libraries for all five sequenced AML samples. Indexed AML samples were subsequently pooled at equal molar concentrations and the resulting pool sequenced on a single MinION device run (**Methods**).

Pooled AML Nanopore run yielded slightly fewer number of sequenced molecules compared to previous short nanopore runs; ∼3.2 million reads. Regardless, the sequencing data was of high quality (93% passed filter and 11.15 Mean Q-score) enabling 91% of the sequenced molecules to be assigned to a sample barcode (average of ∼600,000 reads per sample, 18% of total reads per sample) (**S4 Fig**). When processed for CNA analysis, the multiplex data generated largely similar genome-wide copy number profiles across all samples compared to matching non-multiplex data, with Pearson correlation coefficients above 0.95 (**Fig 3A, B, and S4B and C Fig)**. Identified CNAs, which were similarly detected in non-multiplex nanopore and Illumina sequencing data, included both deletions as well as gains, either as whole chromosome or interstitial, focal alterations, with events targeting important AML genomic regions such as 5q, 7q and 17p (**Fig 3B and S5A Fig)**. Events were confirmed orthogonally using DNA-FISH experiments (**Fig 3C)**.

**Fig3.**
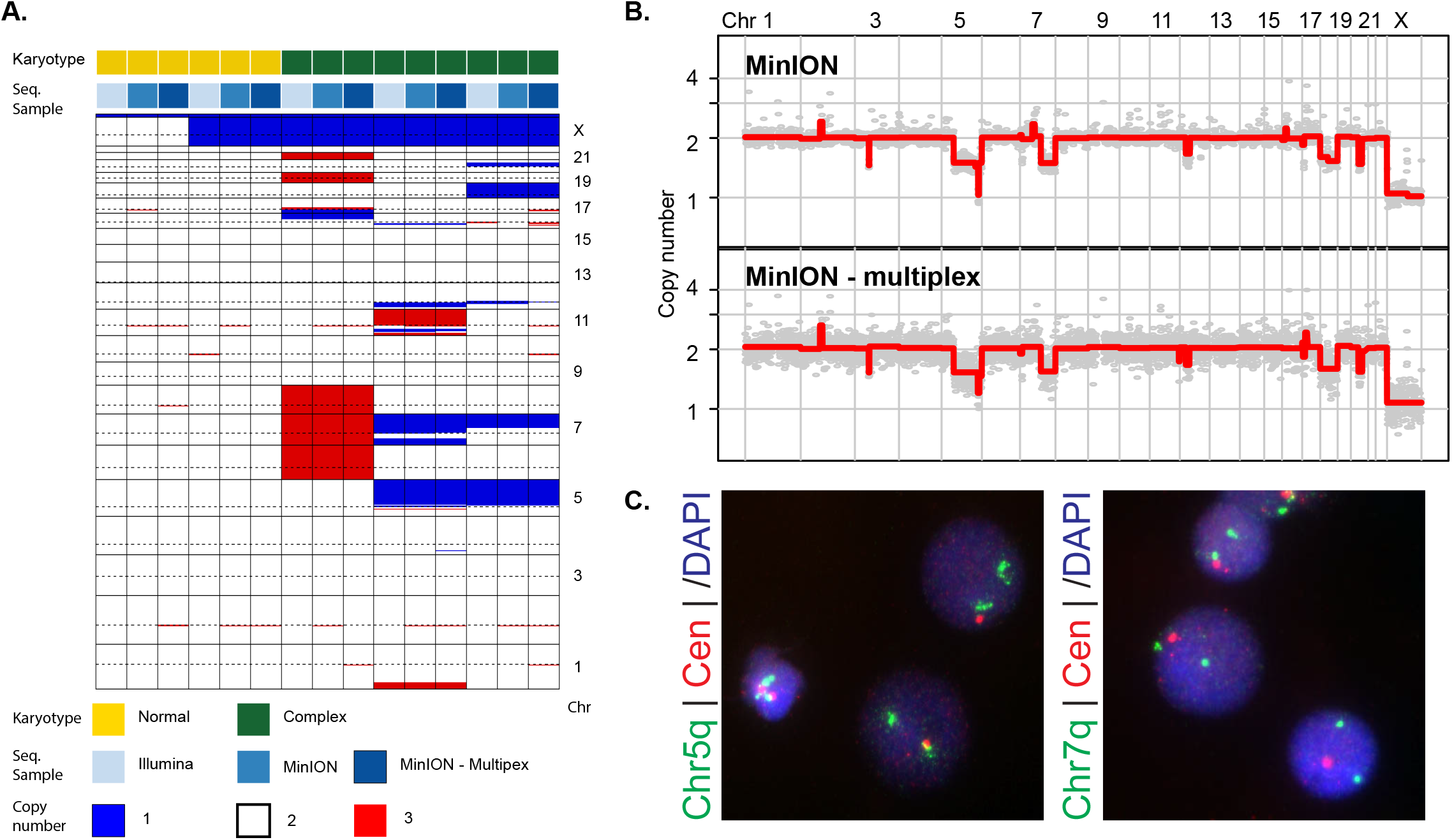
Increased read counts via short molecule sequencing enables accurate, multiplex profiling on a MinION device. (A) Heatmap illustration of copy number profiles of all profiled AML samples (n=5) using Illumina, MinION, and Multiplex MinION sequencing. Annotations regarding karyotype status of leukemic sample (normal vs. complex) and sequencing modality are denoted on bars on top of the heatmap. Heatmap color and bar color codes are provided below the heatmap. (B) Genome-wide copy number profiles from a complex karyotype AML sample inferred from short molecule nanopore sequencing in multiplex (lower panel) and non-multiplex mode (upper-panel). (C) DNA-FISH based validation of the identified deletions of the long arm of chromosomes 5 and 7 (Chr5q and Chr7q, respectively). Probe at 5p15.2 and 7q31 as well as centromeric probes (for internal control) were used. Color codes of probes are illustrated.

Thus, the increase in return in terms of the number of high quality sequencing molecules afforded by loading short DNA fragments on a nanopore device facilitates multiplex sequencing and accurate inference of copy number information in an affordable and versatile manner.

### Short molecule sequencing using a Flongle adaptor yields high read counts and allows cheap portable sequencing

The MinION device has generally been known as a single-use nanopore sequencing device. Recently, Oxford Nanopore introduced the Flongle adaptor; a low-cost replaceable apparatus that utilizes similar chemistry in producing single molecule sequence data. This, in effect, makes the MinION device a repeat-use sequencing device. High read-count recovery via short molecule sequencing using a Flongle device theoretically can enable the retrieval of clinically actionable CNA data (i.e. karyotypes and CNA information) in settings outside large health care centers as well as countries with poorly developed health care infrastructure. To assess the feasibility of such applications, we generated short molecule nanopore libraries for a normal karyotype AML case and a complex karyotype AML case and using a Flongle adaptor for each, sequenced both. Short-read Illumina sequencing was also performed on DNA available for both cases.

Similar to short molecule sequencing on a non-Flongle MinION unit, Flongle based sequencing continued to return relatively high, short molecule sequence information over the first 12 hours of the sequencing run with diminishing returns thereafter (**Fig 4A and S5A)**. Short read Flongle runs yielded much lower numbers of sequenced molecules in terms of absolute counts; ∼600k and 300k for Flongle runs compared to ∼ 4 – 6 million MinION runs (**S5B Fig)**. However, when analyzing for number of sequenced molecules while normalizing for the number of sequencing channels (Flongle contains 126 channels compared to the standard 512 for a non-Flongle MinION unit), the data illustrate that Flongle short molecule sequencing returns almost one order of magnitude more data than a long molecule run (**Fig 4B)**. The pattern of sequenced molecule return was similarly associated with a shorter vacancy time compared to long molecule sequencing, although vacancy time was longer than that observed with non-Flongle MinION Short-1ME runs (**S5C Fig)**. While Flongle runs yielded lower statistics for Passed and Mean Q-Score (**S5B Fig)**, approximately 176,000 and 327,000 reads (83 and 86 % of total pass filter reads) were uniquely mapped, enough to process for copy number analysis. Analysis of the Flongle nanopore data revealed that while one sample exhibited a normal copy number profile, the other sample displayed a pattern of deletions on chromosomes 1, 8, 9, and 11 resulting in complex karyotype annotations, consistent with matching cytogenetic data (**Fig 4C and D)**.

**Fig4.**
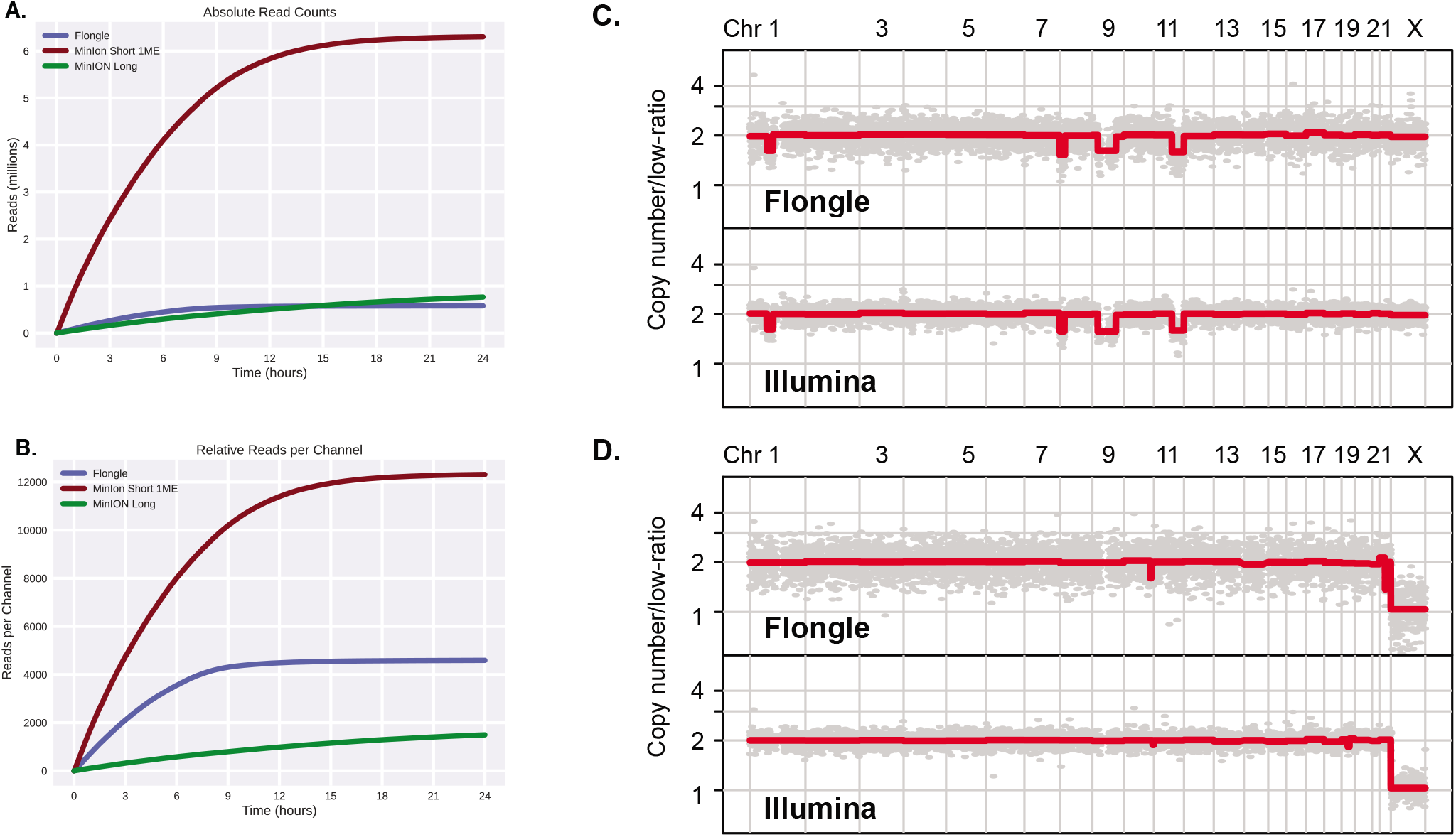
Short molecule sequencing using a Flongle adaptor enables high read counts, accurate copy number data, and cheap, portable sequencing. (A) Absolute number of sequenced molecules for a Flongle run compared to other nanopore runs. Note, Short 1ME and Long are as depicted in Fig1A. (B) Normalized number of sequenced molecules per channel for each nanopore run (126 channels for Flongle and 512 channels for MinION). (C and D) Genome-wide copy number profiles of a complex karyotype (C) and a normal karyotype (D) AML case infered from short molecule nanopore sequencing (top-panel) and short read Illumina sequencing (lower panel), respectively.

Together, the data shows that the Flongle adaptor, when coupled with short molecule sequencing, is capable of facilitating sequencing applications that require a discrete number of sequenced molecules (ex: copy number profiling at specific resolutions). Furthermore, given the early stage nature of the Flongle in terms of development, we expect further modifications/improvements to yield more data within the context of short molecule sequencing in future iterations, and in doing so, allow higher resolution copy number analysis.

## DISCUSSION

Nanopore single molecule sequencing offers unique advantages compared to Illumina short read sequencing including real-time sequencing, portable sequencing, and the potential to sequence long molecules of DNA. Numerous studies have now shown the utility of each of the aforementioned advantages in the rapid re-identification of human samples, facilitating fast point-of-care diagnostics for infectious diseases, the first complete, telomere-to-telomere sequence of a human chromosome and several other applications (Zaaijer et al. 2017; Meredith et al. 2020; Miga et al. 2020; Alonge et al. 2020). However, an important limitation of nanopore sequencing continues to be the relatively low number of sequencing reads returned per sequencing run, regardless of instrument/device (i.e. MinION, GridION, or PromethION). This presents challenges for nanopore sequencing utility in “counting” applications such as allelic genotyping/single nucleotide discovery and genome copy number inference, applications that require high read counts, at specified loci or distributed over a given genomes, respectively. This is important as methods for the identification of CNAs, particularly hematological malignancies, have long been considered important tests/procedures to perform in the course of the clinical management of patients (Döhner et al. 2017). Furthermore, increasing evidence has emerged regarding the importance of CNAs in more common, solid malignancies such as breast and lung cancer (Curtis et al. 2012; Sansregret, Vanhaesebroeck, and Swanton 2018).

Here, by (1) making a simple optimization with regards to molecule length when factoring DNA molecule kinetics during translocation through a nanopore and (2) applying an elementary strategy of loading short DNA molecules on a MinION device, we devise and validate a simple yet effective solution: short molecule nanopore sequencing. Loading short molecules of DNA (median length of 0.5kb) on a MinION device consistently resulted in a 4 to 6-fold increase in the return of sequenced single molecules. Importantly, the resulting CNAs from the short molecule data were of similar quality to data obtained via standard short-read Illumina sequencing and with much higher resolution than clinical karyotyping.

The short molecule nanopore sequencing approach presented here has broad applications in research and clinical practice including, but not limited to, global DNA modification profiling via shotgun DNA sequencing as well as multiplex cancer gene amplicon sequencing. Most importantly, we show here that high molecule return on a nanopore can enable multiplex sequencing, that when coupled with molecular barcoding, vastly decreases the costs and time associated with the retrieval of clinically important information, such as karyotype information in AML disease. Additionally, in relation to sequence associated costs and potential clinical utility, we present results of short molecule nanopore sequencing within the context of the Flongle adaptor, providing further evidence for the potential for cost effective, point-of-care diagnostics using nanopore sequencing technology in cancer.

While others have devised innovative approaches using nanopore sequencing to address genomic applications that require high molecule/read counts (Prabakar et al. 2019), our approach is much simpler, especially for barcoded multiplexed samples. Hence, we surmise that short molecule nanopore sequencing is more likely to gain broad applicability and equally important, empower further technological developments in the rapidly evolving field of single-molecule sequencing. For example, we have recently reported on the development of an algorithm termed UNCALLED for the targeted enrichment of genomic sequences based on analysis of raw electrical nanopore signals and ejection of unwanted molecules from any given pore (Kovaka et al. 2020). Coupling the methods used in short molecule sequencing with targeted enrichment of specific loci could enable the retrieval of the entire repertoire of germline and somatic alterations found in a cancer, including single nucleotide variants, insertions and deletions as well as balanced structural variants all from a single sequencing run. Indeed, our early results point towards the feasibility of such an approach.

In conclusion, we show that short molecule nanopore sequencing facilitates increased sequence molecule return, enabling applications that require sequencing counts, such as CNA detection, that are inaccessible with standard, current nanopore practices (i.e. long molecule sequencing). We show that the data is high quality and when applied to address a biomedical question, such as retrieving clinically meaningful copy number information, returns accurate data at low costs and much flexibility in operation. We posit that short molecule nanopore sequencing will have broad applications, in DNA as well as RNA sequencing, that extend beyond genome copy number quantification.

## ACKNOWLEDGEMENTS

We would like to thank the patients that donated their samples for this study. This work was supported, in part, by the National Institutes of Health (5R50CA243890 to S.G. and R01 CA190261 to SWL) and the US National Science Foundation (DBI-1350041 to M.C.S.). TB is supported by the William C. and Joyce C. O’Neil Charitable Trust, Memorial Sloan Kettering Single Cell Sequencing Initiative. SWL is an investigator in the Howard Hughes Medical Institute and the Geoffrey Beene Chair for Cancer Biology.

## SUPPLEMENTAL INFORMATION

**S1 Fig**. Short DNA molecule sequencing on a MinION device yields a high number of sequencing reads.

**S2 Fig**. MinION short molecule sequencing data returns highly accurate genome-wide copy number information.

**S3 Fig**. Short molecule sequencing on a MinION device reproducibly yields accurate, high resolution copy number information.

**S4 Fig**. Multiplex short molecule sequencing on a MinION device allows accurate and cheap inference of CNAs.

**S5 Fig**. Short molecule sequencing using a Flongle adaptor returns high read counts and allows accurate copy number inference.

