## Supplemental Figure 1 for "High resolution copy number inference in cancer using short-molecule nanopore sequencing"

### Supplementary Figure 1

**A.**

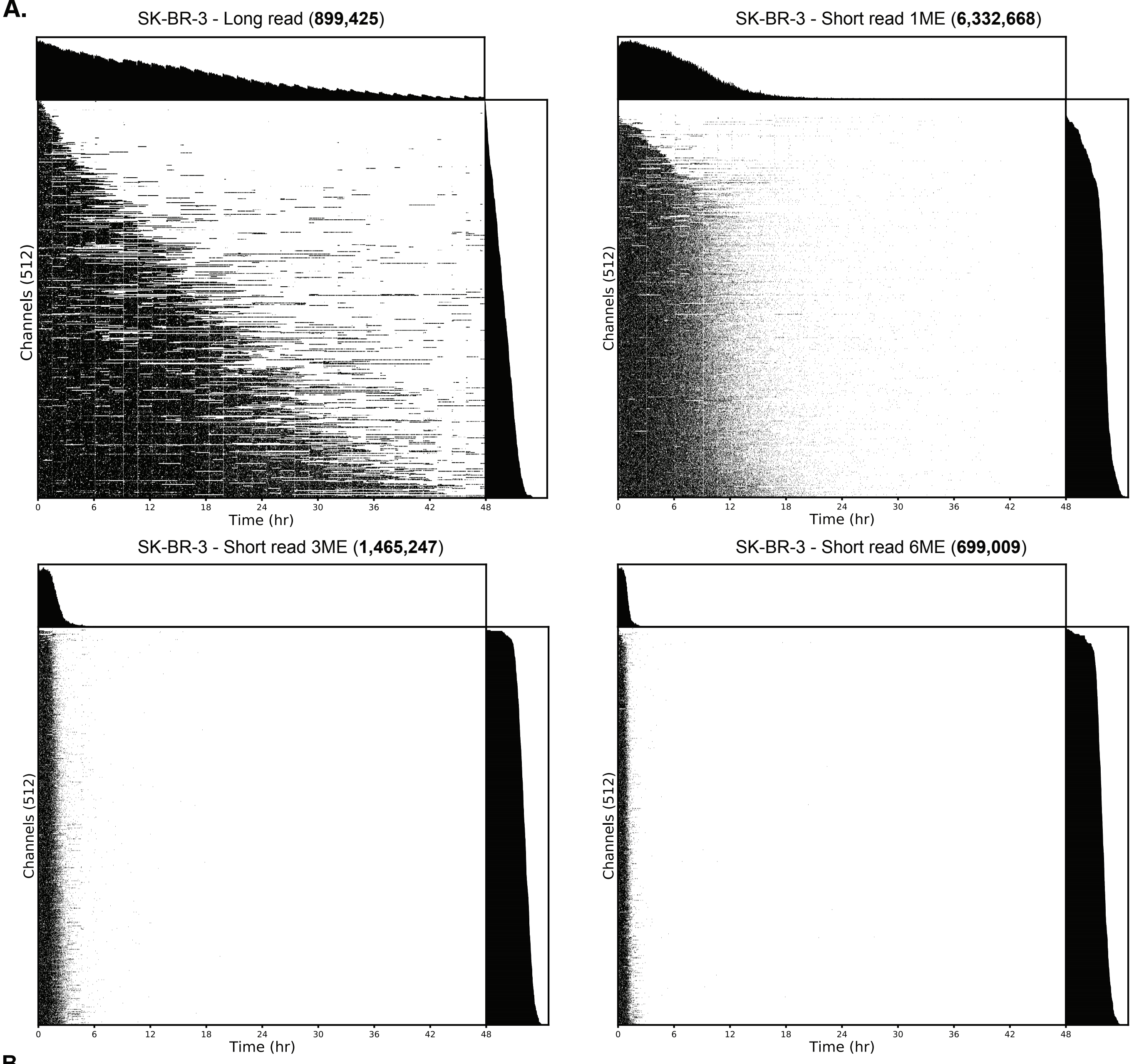


**B.**

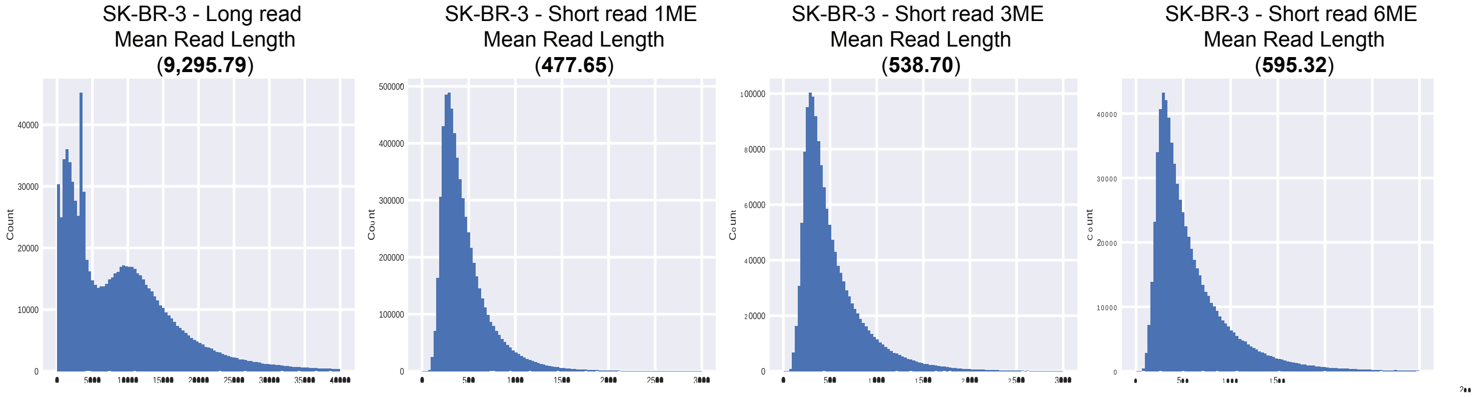


**S1 Fig. Short DNA molecule sequencing on a MinION device yields high number of sequencing reads.**
**(A)** Relationship between nanopore occupancy and read count yields on a MinION device using different SK-BR-3 sequencing library preparations and loading conditions. **(B)** Read length distrubition of the different SK-BR-3 sequencing library preprations and loading conditions as in (A).
