## Supplemental Figure 2 for "High resolution copy number inference in cancer using short-molecule nanopore sequencing"

### Supplementary Figure 2

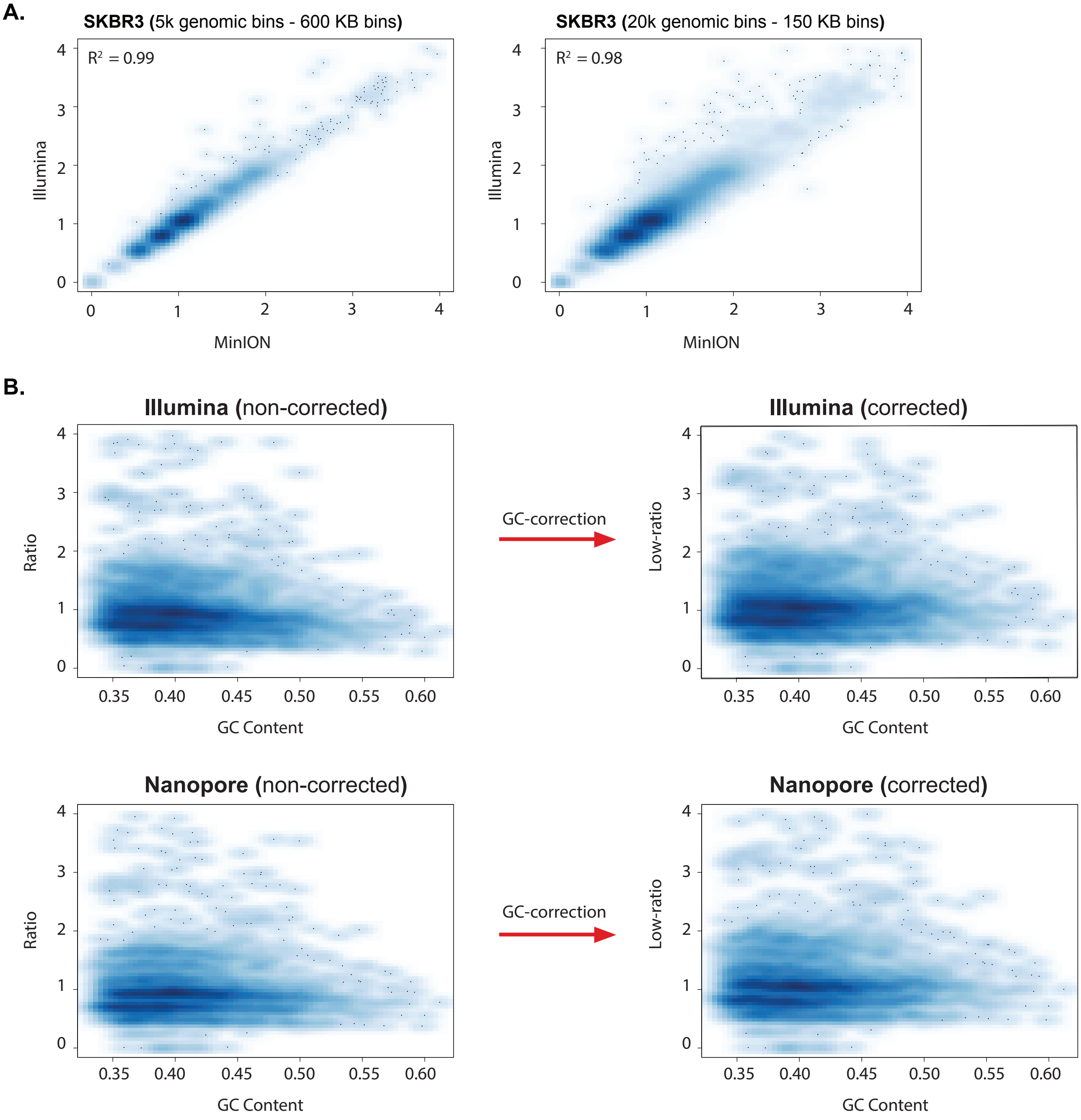


**S2 Fig. MinION short molecule sequencing data returns highly accurate genome-wide copy number information.** Scatter density correlation plots of normalized read count values from Illumina and MinION short read (Short 1ME) data. Scatter density plots are illustrated for two resolutions; 5k and 20k (i.e. 5000 genomics bins and 20,000 genomics bins respectively). Pearson correlation values for the analysis at each resolution are provided. **(B)** Scatter densityplots illustrating GC bias in the SKBR3 Illumina and Short-1ME nanopore data (left panels) and scatter density plots of corrected data using LOWESS smoothing (right panels).
