## Supplemental Figure 3 for "High resolution copy number inference in cancer using short-molecule nanopore sequencing"

### Supplementary Figure 3

**A.**

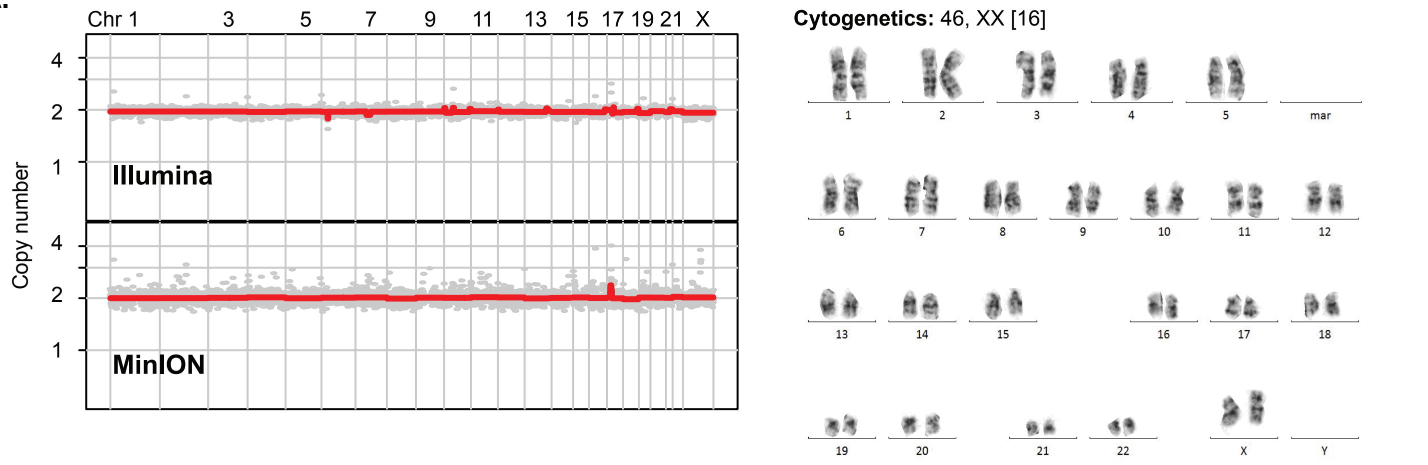


**B.**

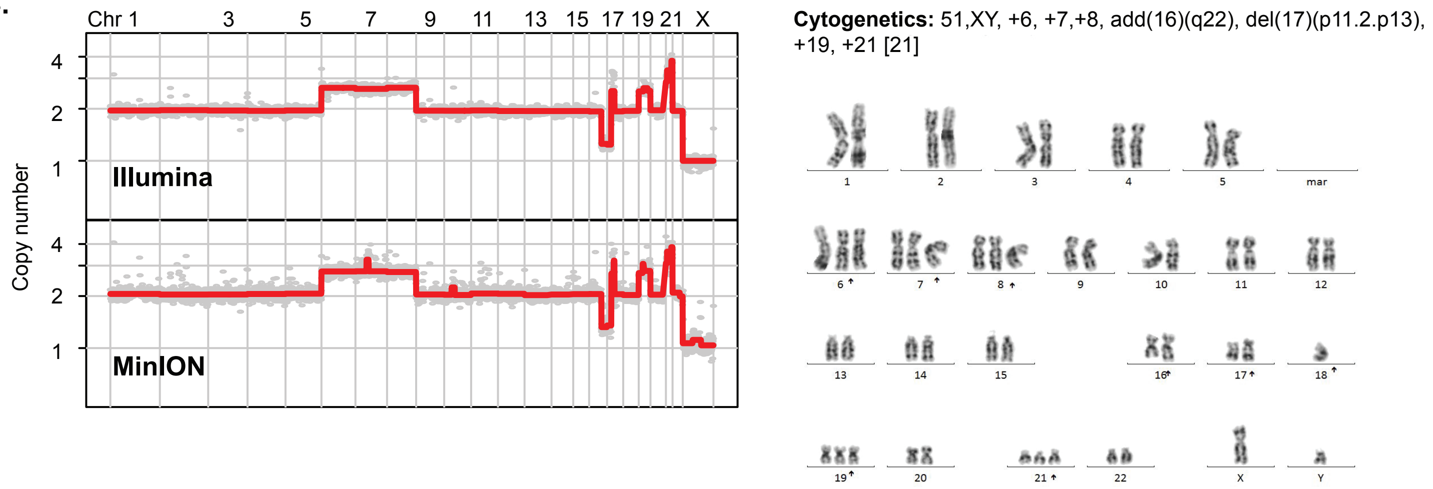


**C.**

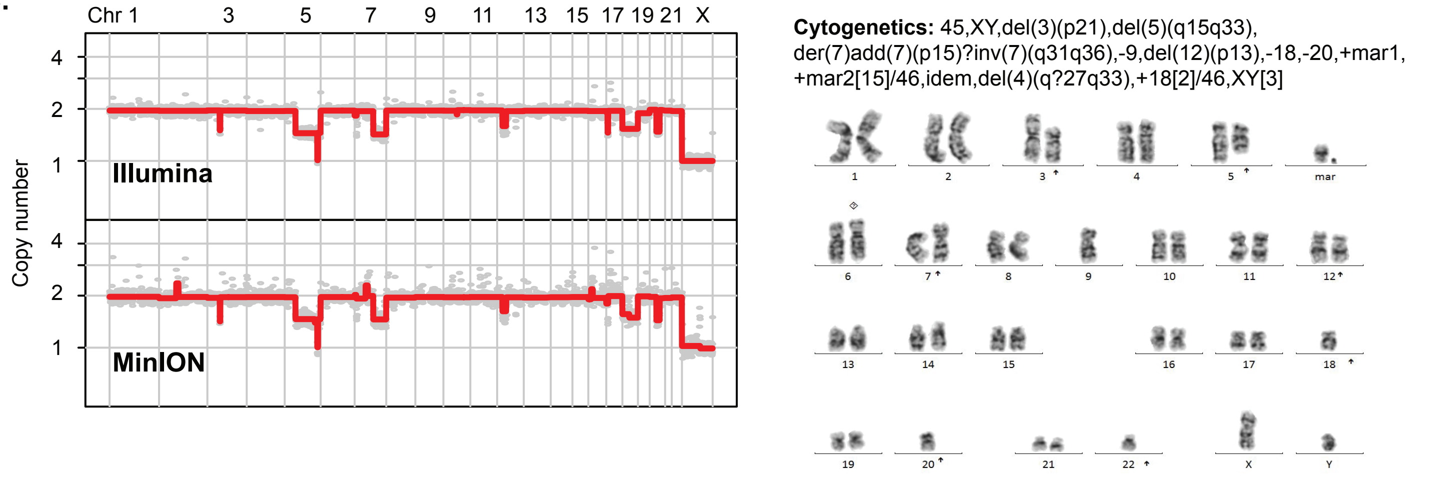


**S3 Fig. Short molecule sequencing on a MinION device reproducibly yields accurate, high resolution copy number information. (A-C)** Left panels **-** Genome-wide copy number profiles of ormal and complex karyotype samples sequenced using short read/molecule nanopore sequencing on a MinION (lower panel) and an Illumina (upper-panel) device.
Right panels - matching chromosome spread, microscopy images of cytogenetically analyzed AML samples. Cytogenetic annotations are provided on to of the microscopic images.
