## Supplemental Figure 4 for "High resolution copy number inference in cancer using short-molecule nanopore sequencing"

### Supplementary Figure 4

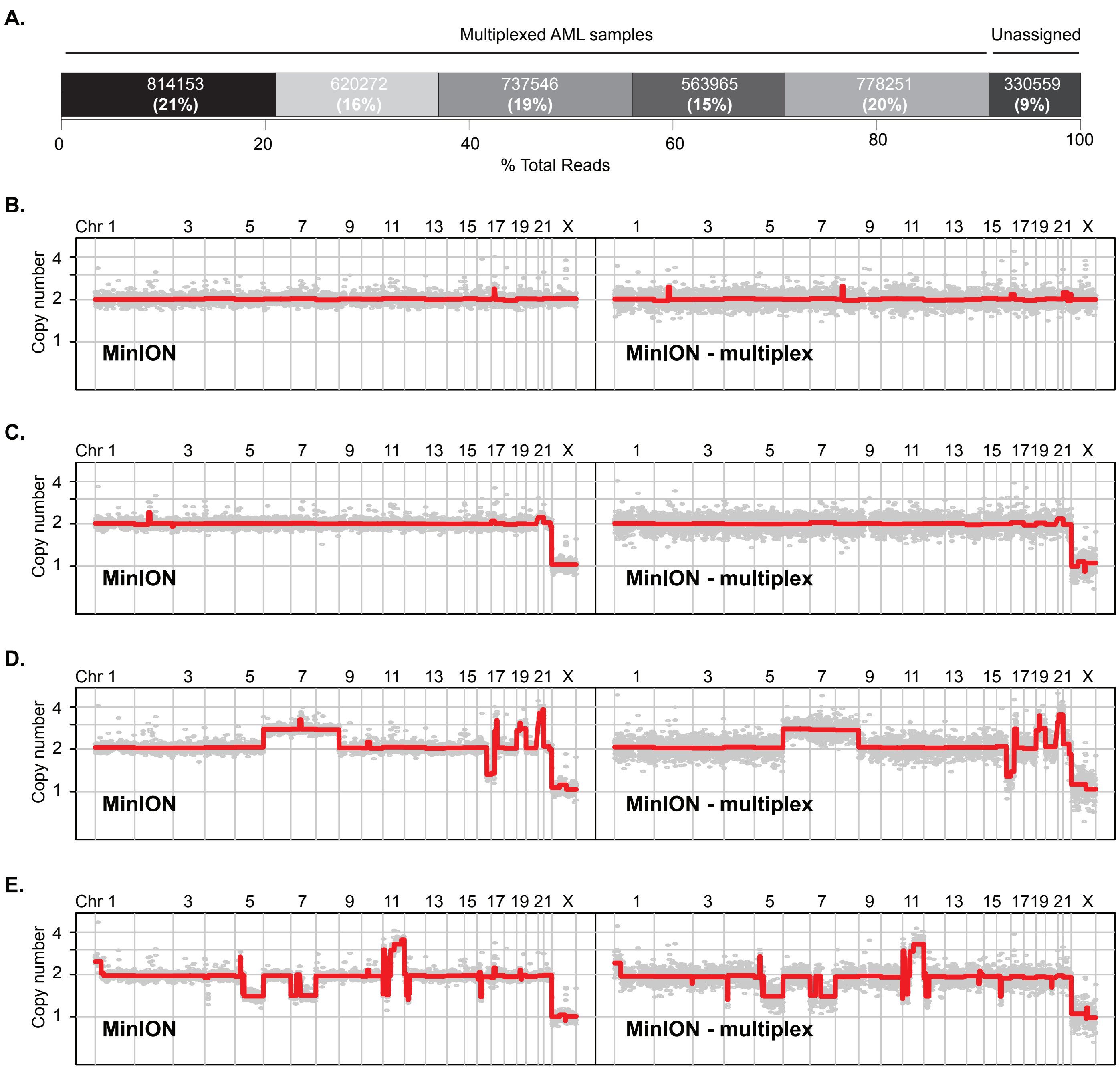


**S4 Fig. Multiplex short molecule sequencing on a MinION device allow accurate and cheap inference of CNAs. (A)** Bar plot quantification of de-multiplexed sequencing reads and percentages per pooled, barcoded AML samples plus unassigned reads. **(B-E)** Genome-wide copy number profile of a two normal karyotype (B,C) and two complex karyotype (D, E) samples sequenced on a MinION device in non-multiplex (left panel) and multiplex (right panel) fashion.
