## Supplemental Figure 5 for "High resolution copy number inference in cancer using short-molecule nanopore sequencing"

### Supplementary Figure 5.

A.

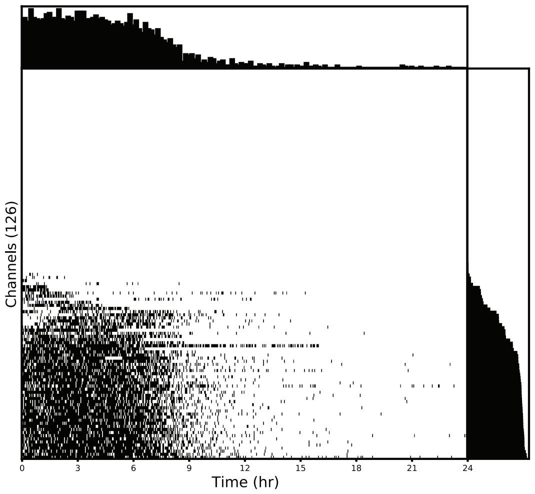


C.

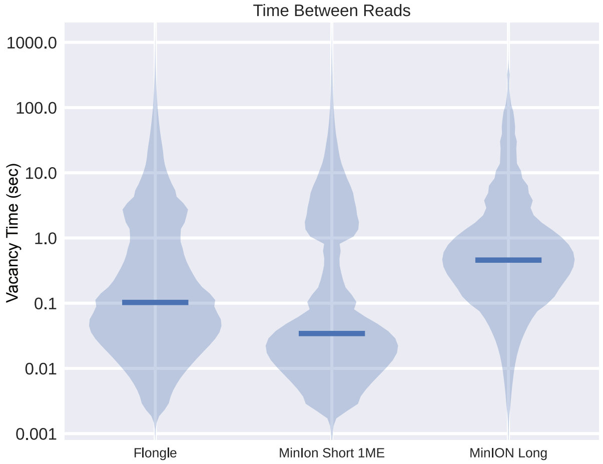


B.

| Sample/ ID | Run.Type | Read Count | Yield (Mbp) | Passed (%) | Q-Score Mean | Read- Length Mean | Read N50 | Minutes to 250k reads |
|---|---|---|---|---|---|---|---|---|
| AML2 | MinION | 6,263,490 | 2,886.35 | 82.82 | 8.95 | 460.82 | 519 | 15.53 |
| AML3 | MinION | 3,935,726 | 1,753.57 | 86.68 | 9.29 | 445.55 | 488 | 15.58 |
| AML6 | **Flongle** | 578,420 | 257.10 | 65.36 | 8.26 | 444.48 | 479 | 167.70 |
| AML7 | **Flongle** | 296,062 | 128.04 | 71.87 | 8.52 | 432.49 | 473 | 294.52 |

**S5 Fig. Short molecule sequencing using a Flongle adaptor returns high read counts and allows accurate copy number inference. (A)** Channel activity throughout the Flongle run - see Fig1 for detailed explanation of figure. **(B).** Flongle short molecule sequencing metrics compared to MinION runs. **(C)** Distributions of vacancy time (i.e. time between when one molecule finishes and one starts) for Flongle run, compared to the long and short 1ME MinION runs.
